# Age dependent effects of protein restriction on dopamine release

**DOI:** 10.1101/2020.04.09.033985

**Authors:** Fabien Naneix, Kate Z. Peters, Andrew M. J. Young, James E. McCutcheon

## Abstract

Despite the essential role of protein intake for health and development, very little is known about the impact of protein restriction on neurobiological functions, especially at different stages of the lifespan. The dopamine system is a central actor in the integration of food-related processes and is influenced by physiological state and food-related signals. Moreover, it is highly sensitive to dietary effects during early life periods such as adolescence due to its late maturation. In the present study, we investigated the impact of protein restriction either during adolescence or adulthood on the function of the mesolimbic (nucleus accumbens) and nigrostriatal (dorsal striatum) dopamine pathways using fast-scan cyclic voltammetry in rat brain slices. In the nucleus accumbens, protein restriction in adults increased dopamine release in response to low and high frequency trains of stimulation (1-20 Hz). By contrast, protein restriction performed at adolescence decreased nucleus accumbens dopamine release. In the dorsal striatum, protein restriction has no impact on dopamine release when performed at adulthood but in adolescent rats we observed frequency-dependent increases in stimulated dopamine release. Taken together, our results highlight the sensitivity of the different dopamine pathways to the effect of protein restriction, as well as their vulnerability to deleterious diet effects at different life stages.

## INTRODUCTION

The regulation of food intake in an ever-changing environment is a central survival process. Healthy diet requires a balanced intake of the three main macronutrients (carbohydrate, fat, and protein) [1]. Protein intake is especially important as amino acids are essential for many biological functions (growth and maintenance, synthesis of nucleic acids and hormones, immune response, cellular repair) and many amino acids cannot be synthesized *de novo*. In humans, protein deficiency and a low protein diet are associated with muscle wasting, stunted growth and increased vulnerability to infections, but may also, to some extent, contribute to obesity by generally increasing appetite [2–4]. Furthermore, protein deficiency and severe protein malnutrition are especially detrimental during development and early life when demand is highest [5–7]. Numerous species, including humans and rodents, regulate their food intake and food-related behaviors to avoid protein deficiency [8–13]. Increasing evidence implicates broad hypothalamic and limbic circuits in the regulation of protein appetite [10,13–15]. However, the impact of protein imbalance (high or low protein diet) on the function of these neurobiological circuits remains undescribed, especially when protein deficiency occurs during a critical period of early development.

The dopamine system plays a central role in food-seeking behaviors, food preference, and in the motivation to eat [16–19]. Recent data show that dopamine neurons integrate current physiological state (*i*.*e*. hunger, nutrient deficiency) to guide food-seeking behaviors [20–23]. Dopamine neurons are especially sensitive to the nutrient content of ingested food [24–28], through gut-to-brain axis [29,30] and peripheral feeding hormones [31–35]. Furthermore, exposure to specific diets, such as high-carbohydrate and/or high-fat, impacts dopamine signaling within the nucleus accumbens (NAc) and the dorsal striatum [36–40]. However, the impact of low protein diet on the function of dopamine circuits is still largely unexplored.

Early life periods like childhood and adolescence are periods of particular vulnerability to the deleterious impact of various diets on corticolimbic circuits and reward-related processes [41–47]. Interestingly, the dopamine system undergoes delayed maturation taking place during adolescence making it vulnerable to external insults [47–54]. The impact of prolonged inadequate protein consumption on dopamine signaling remains unknown but may be exacerbated during adolescence when protein demand is increased to support rapid growth 65 [55].

Here, we investigated the impact of protein restriction either during adolescence or adulthood on the function of the mesolimbic (NAc) and nigrostriatal (dorsal striatum) dopamine pathways using fast-scan cyclic voltammetry (FSCV) in rat brain slices. We found that protein restriction induced opposite effects on NAc dopamine release depending on age, with restriction increasing dopamine release in adults but decreasing it in adolescents. In the dorsal striatum, however, dopamine function following protein restriction was increased only in adolescents and not adults.

## MATERIAL AND METHODS

### Subjects

Male Sprague Dawley rats (Charles River Laboratories) were received either at weaning (approximately P21, 50-70 g) for adolescent groups (n = 13) or at adulthood (P60, 200-250 g) for Adult groups (n = 15). Rats were housed in groups of 2-3 in individually ventilated cages (46.2 x 40.3 x 40.4 cm), in a temperature (21 ± 2°C) and humidity (40-50%) controlled environment with a 12 h light/dark cycle (lights on at 7:00 AM) and with food and water available *ab libitum*. All testing and tissue harvesting occurred in the light phase. Procedures were performed in accordance with the Animals (Scientific Procedures) Act 1986 and carried out under Project License PFACC16E2.

### Diets

All rats were initially maintained on standard laboratory chow diet (Teklad global #2918, Envigo) containing 18% protein. One week after arrival rats either continued on standard laboratory chow diet (Non Restricted group; Adolescents-NR n = 6, Adults-NR n = 8) or were switched to a modified AIN-93G diet containing 5% protein from casein (#D15100602, Research Diets; Protein Restricted group: Adolescents-PR n = 7, Adults-PR n = 7; **Supplemental Table 1**) [11]. Rats had *ad libitum* access to their assigned diet. Protein restriction was maintained for 12 to 14 days either during adolescence (from P28 to P42) or during adulthood (> P70). Body weight and food intake were collected daily throughout the experiments. Tissue was collected for voltammetry recordings immediately after this period (**Figure 1A**).

**Figure 1.**
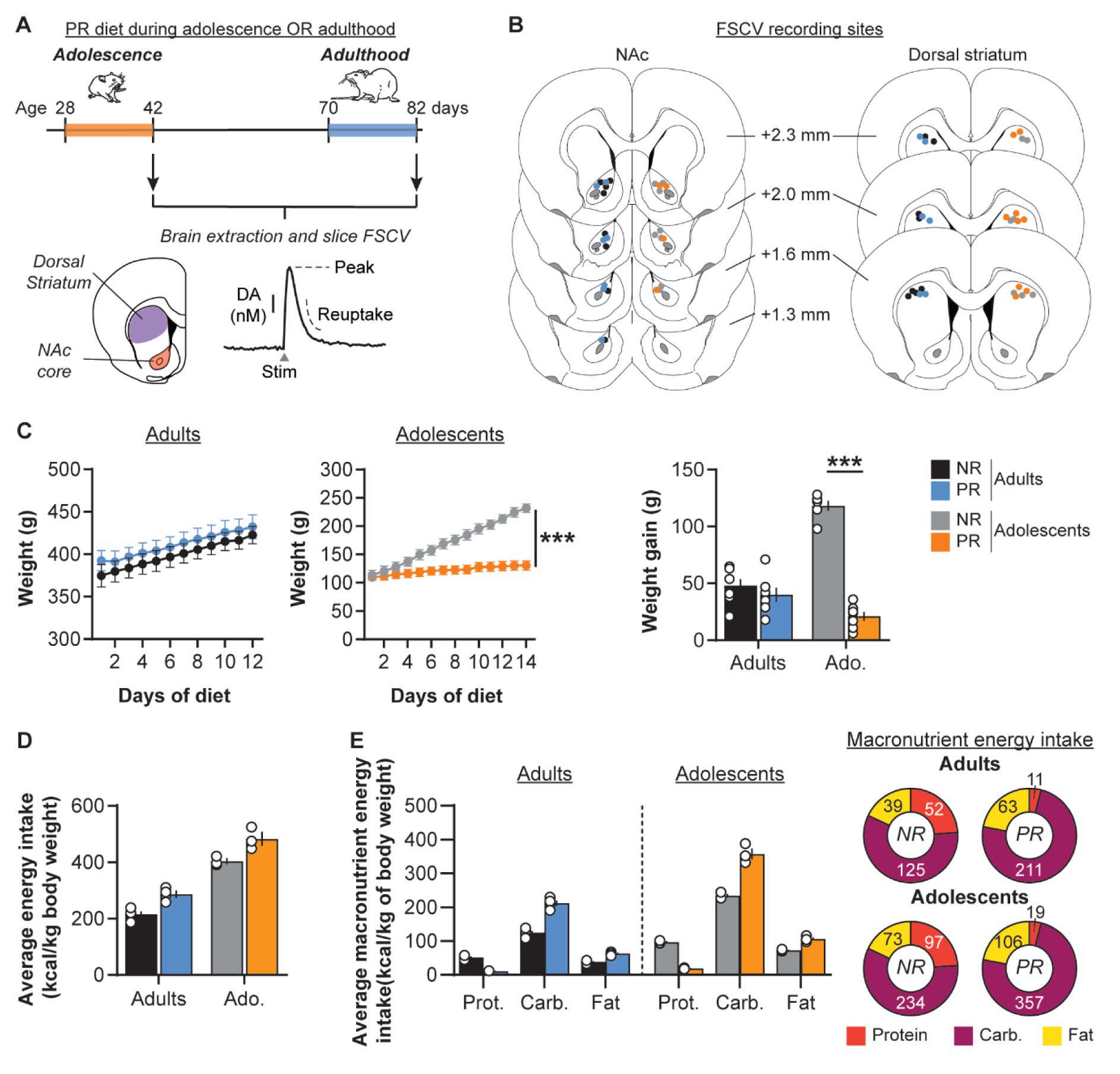
(A) Schematic representation of the experimental design. Rats had access to either control chow diet (18% protein; non-restricted, NR) or protein-restricted diet (5% protein; PR) during either adolescence (post-natal day 28 to 42) or adulthood (post-natal-day 70 to 82). At the end of the diet exposure, brains were extracted to perform FSCV recordings of electrically evoked dopamine (DA) release in the NAc and dorsal striatum. (B) Coronal brain sections (modified from [83]) representing the recording sites in the NAc core (*left*) and the dorsal striatum (*right*) for NR and PR groups. Numbers indicate the distance from Bregma. (C) Protein restriction during adolescence but not during adulthood altered weight (*left and middle*) and weight gain (*right*). (D) Daiiy energy intake (in kcal/kg of body weight) is higher in adolescent rats compared to adults, and in PR groups compared to control NR groups. (E) Macronutrient breakdown for daily energy intake during the diet exposure (in kcal/kg of body weight). Pie charts represent energy intake from each macronutrient for each diet group (Protein: red; Carbohydrate: purple; Fat: yellow). Adults-NR, n = 8, black symbols; Adults-PR, n = 7, blue symbols; Adolescents-NR, n = 6, grey symbols; Adolescents-PR, n = 7, orange symbols. Data are mean ± SEM and circles show individual (*e*.*g*. rats for C and cages for D) data points. *** *p* < 0.001 Diet effect (two-way ANOVA followed by Sidak’s *post hoc* tests).

### Slice preparation

Rats were deeply anesthetized with chloral hydrate (400 mg/kg i.p., Sigma-Aldrich), decapitated, and brains were removed and transferred to ice cold artificial cerebrospinal fluid (aCSF) containing in mM: 126 NaCl, 10 glucose, 26 NaHCO_3_, 2.5 KCl, 2.4 CaCl_2_, 2 MgCl_2_, 1.4 NaH_2_PO_4_. Acute 300 µm thick coronal slices, containing both the NAc and the dorsal striatum were prepared in ice-cold aCSF buffer using a vibratome (Leica VT1200S). Slices were kept at room temperature (20-22°C) in aCSF saturated with 95% O_2_ and 5% CO_2_ for at least 1 h before the start of recordings.

### Fast scan cyclic voltammetry recordings

Unilateral slices were transferred to the recording chamber and superfused at 2 ml/min with aCSF saturated with 95% O_2_ and 5% CO_2_ at 30°C. Slices were allowed to equilibrate for 30 min prior to recordings. A twisted stainless steel bipolar stimulating electrode (MS303T/2-B/SPC, P1 Technologies) was placed at the surface of the slice within the NAc core or the dorsal striatum (**Figure 1B**). A homemade glass capillary carbon-fiber microelectrode (tip length 50-100 µm) was positioned in the slice approximately 100 µm beneath the tissue surface and 100-200 µm from the stimulating electrode [56,57]. For FSCV recordings, a triangular voltage waveform was applied (−0.4 to +1.3 V and back versus an Ag/AgCl reference electrode; 400 V/s) using a custom-built headstage circuit (University of Washington Electronics and Materials Engineering Shop, Seattle, WA) and TarHeel voltammetry software (Chapel Hill, University of North Carolina [58]). The waveform was initially applied at 60 Hz for 10 min, to condition the electrode outside of the tissue, and then applied at 10 Hz while all experiments were being conducted. Dopamine release was evoked by monopolar stimulation pulses (0.7 mA, 0.2 ms) [59]. Electrical stimulations were repeated at 3 min intervals to ensure consistent release. Stimuli were either single pulses (1 p) or trains of five pulses (5 p) at frequencies ranging from ‘tonic’ (1, 5 or 10 Hz) to ‘phasic’ burst frequencies (20 Hz) of dopamine neurons reported *ex vivo* and *in vivo* [60–62]. Each stimulation was repeated 3 times in pseudo-random order and averaged to obtain the individual value for this frequency. Each slice yielded an individual recording site.

Extracellular dopamine levels ([DA]_o_) were confirmed by examining current-voltage plots showing oxidation (approximately +0.6 V) and reduction (approximately −0.2 V) peaks using TarHeel software. Background (non-Faradaic) current was measured for 1 s between 4-5 seconds before the stimulation and subtracted from the signal. Dopamine currents (in nA) were then converted to dopamine concentration (in nM) using the calibration of each electrode against a known standard dopamine concentration. [DA]_o_ peaks were measured following any stimulation artefacts within a +0.2 to +0.5 sec time interval following the start of the stimulation, as previously described [63]. As the electrical stimulations used varied in length and frequency, we also quantified DA release by using the area under the curve of [DA]_o_ (AUC) following the start of the stimulation. Recording electrodes were calibrated after use using 1-2 µM dopamine solution in a flow cell system [64] and in the recording chamber.

### Statistical analysis

Weight and food intake measures were analyzed using three- or two-way repeated measures ANOVAs with Diet (Non Restricted NR, Protein Restricted PR) and Age (Adults, Adolescents) as between factors and Day or Macronutrient (Protein, Carbohydrate, Fat) as within factors. As rats were group-housed, food intake data were collected by cage, divided by the number of rats in the cage, normalized by kg of body weight and expressed as energy intake (kcal/kg of body weight). Energy intake was also analysed as macronutrient breakdown.

For single pulse stimulation, [DA]_o_ AUC and clearance times (T_80_: time for 80% decay from peak amplitude; T_20_: time for 20% decay from peak amplitude; Half-life: time for 50% decay from peak amplitude) were analyzed using two-way ANOVAs with Age (Adults, Adolescents) and Diet (NR, PR) as between-subject factors. [DA]_o_ AUC in response to single pulses were plotted as cumulative probability and compared using Kolmogorov-Smirnov test. [DA]_o_ AUC from frequency-response curves were analyzed using three-way and two-way repeated measures ANOVA using Age (Adults, Adolescents) and Diet (NR, PR) as between-subject factors and Frequency (1, 5, 10, 20 Hz) as within-subject factor. 5 p / 1 p [DA]_o_ ratios were calculated by dividing the average [DA]_o_ peak value at 20 Hz by the average [DA]_o_ peak value at 1 Hz for the same recording site, and were analyzed using two-way ANOVA with Age (Adults, Adolescents) and Diet (NR, PR) as between-subject factors. Sidak and Dunnett *post hoc* tests were performed when required.

Statistical analyses were conducted using GraphPad Prism 8. All values were expressed as mean ± standard error of the mean (SEM). The alpha risk for the rejection of the null hypothesis was 0.05.

Upon publication, all data analyzed in this paper will be available on Figshare (<u>10.25392/leicester.data.c.5008904</u>).

## RESULTS

### Age dependent impact of protein restriction on weight

We first investigated the impact of protein restriction during either adolescence or adulthood on weight and weight gain (**Figure 1C**). As we previously observed [11], protein restriction at adulthood did not significantly affect rats’ weight (two-way repeated measures ANOVA: Diet, 168 F(1,13) = 0.5, *p* = 0.5; Day, F(11, 143) = 85.5, *p* < 0.001; Diet x Day, F(11, 143)= 0.4, *p* = 1.0). In contrast, protein restriction during adolescence significantly decreased weight gain, relative to control diet (Diet, F(1,11) = 19.8, *p* < 0.001; Day, F(13, 143) = 478.7, *p* < 0.001; Diet x Day, F(13, 143)= 234.0, *p* < 0.001). Both NR and PR adult rats exhibited similar low weight gain (48 g ± 6 and 40 g ± 6, respectively). NR adolescent rats showed substantial weight increases (+118 g ± 4), indicating a normal developmental growth whereas PR rats showed only a modest increase in their weight (+21 g ± 4; two-way ANOVA: Diet, F(1,24) = 23.14, *p* < 0.001; Age, F(1,24) = 97.8, *p* < 0.001; Diet x Age, F(1,24) = 70.3, *p* < 0.001; Sidak’s *post hoc* tests *p* = 0.4 for Adults and *p* < 0.001 for Adolescents), demonstrating that protein restriction in adolescence disrupted normal growth.

Analysis of the average daily food intake for each cage showed that adolescent rats have a higher energy intake than adults (in kcal per kg of body weight; two-way ANOVA: Diet, F(1,11) = 229.8, *p* < 0.001; **Figure 1D**). Moreover, PR groups also exhibited a higher daily energy intake (Diet, F(1,11) = 35.1, *p* < 0.001; Diet x Age, F(1,11) = 0.1, *p* = 0.7). A more detailed analysis of macronutrient breakdown showed that PR groups had an lower energy intake from protein but an increased intake from carbohydrate and fat (3-way repeated measures ANOVA: Diet, F(1,11) = 35.1, *p* < 0.001; Diet x Macronutrient, F(2,22) = 394.7, p < 0.001; all Sidak’s *post hoc* tests *p* < 0.05; **Figure 1E** and **Supplementary Table 2**).

After two weeks of protein restriction, we then assessed the neurobiological impact of this diet on dopamine release in both the NAc and the dorsal striatum using *ex vivo* FSCV in brain slices.

### Age dependent impact of protein restriction on NAc dopamine release

#### Single pulse evoked NAc dopamine release

In the NAc, protein restriction had a different impact on dopamine release evoked by single pulse stimulation depending on the life stage (**Figure 2A-B**; Two-way ANOVA: Age, F(1,24) = 0.1, *p* = 0.7; Diet, F(1,24) = 0.7, *p* = 0.4; Diet x Age, F(1,24) = 17.8, *p* < 0.001). Protein restriction at adulthood induced a significant increase (+167 % ± 49) in NAc dopamine release in response to single pulse stimulation compared to NR control rats (Sidak’s *post hoc* tests *p* < 0.01). In contrast, protein restriction during adolescence significantly decreased Nac dopamine release evoked by single pulse stimulation (**Figure 2A-B**; −44% ± 9; *p* < 0.05). Further analyses confirmed that protein restriction in adulthood significantly changed the distribution of [DA]_o_ AUC values toward the right, demonstrating a greater proportion of large dopamine responses to single pulse, compared to control animals (**Figure 2C**; Kolmogorov-Smirnov test: D(13) = 0.7, *p* < 0.05). In adolescents, protein restriction significantly induced a left-shift of the distribution of [DA]_o_ AUC, confirming a reduced dopamine response (**Figure 2C**; Kolmogorov-Smirnov test: D(11) = 0.8, *p* < 0.05). Importantly, analyses of [DA]_o_ peaks evoked by single pulse confirmed the age-dependent differential effect of the diet (**Figure 2D**; Two-way ANOVA: Age, F(1,24) = 1.4, *p* = 0.2; Diet, F(1,24) = 0.2, *p* = 0.7; Diet x Age, F(1,24) = 7.6, *p* < 0.05), without revealing significant differences in either adults (Sidak’s *post hoc* test *p* = 0.1) or adolescent rats (Sidak’s *post hoc* test *p* = 0.09). To examine whether the diet-induced changes in dopamine release were mediated by differences in dopamine reuptake, we measured the T_80_ clearance time. T_80_ was significantly shorter in adolescent groups compared to adults (**Figure 2E**; Two-way ANOVA: Age, F(1,24) = 6.5, *p* < 0.05). However, protein restriction at adulthood or during adolescence did not seem to significantly change dopamine clearance (Two-way ANOVA: Diet, F(1,24) = 0.1, *p* = 0.7; Diet x Age, F(1,24) = 2.7, *p* = 0.1; see also **Supplementary Figure 1**). Thus, it appears that the robust changes to Nac dopamine release reported as AUC are not driven wholly by either a change to the [DA]_o_ peak amplitude or the time course of dopamine uptake, but likely a combination of both factors.

**Figure 2.**
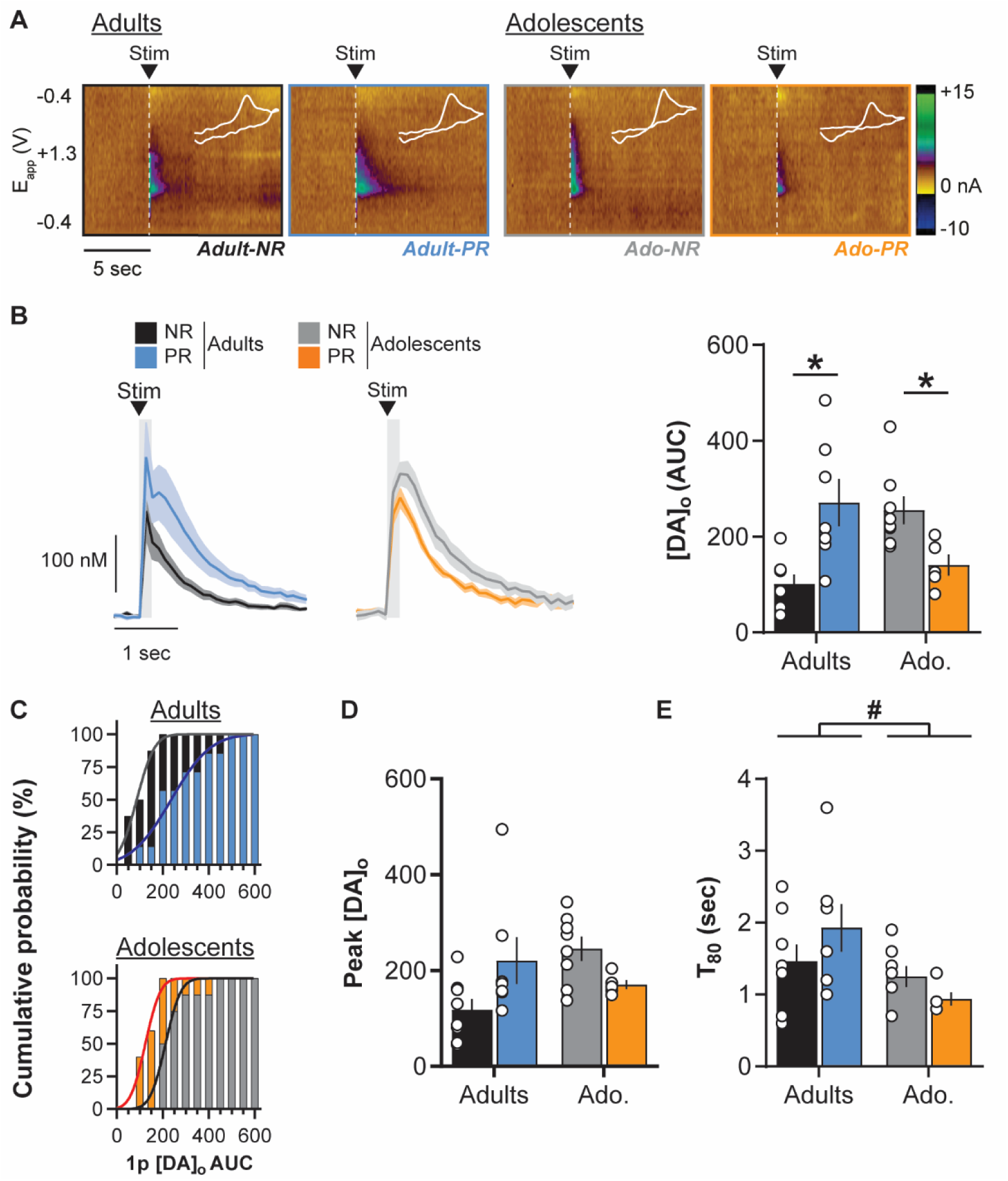
Age-dependent impact of protein restriction on NAc dopamine release evoked by single pulse. (A) Representative FSCV color plots for each diet group (non-restricted, NR; protein-restricted, PR) depicting current changes (color) over time (x-axis; in sec) as a function of the recording electrode holding potential (y-axis; −0.4 to +1.3 V and back) in response to single pulse electrical stimulation (0.7 mA, 0.2 ms; vertical white dashed lines). White line insets represent voltammograms for each color plot. (B) **Left:** NAc [DA]_o_ versus time (in nM; mean ± SEM) in slices from adult and adolescent NR and PR rats, aligned to the single pulse electrical stimulation (black arrow); **Right:** Mean [DA]_o_ release (AUC) evoked by single pulse stimulation in the NAc. (C) Cumulative distribution of single pulse evoked NAc [DA]_o_ AUC in adult (top) and adolescent **(bottom)** groups. (D) Mean [DA]_o_ peak evoked by single pulse stimulation in the NAc. (E) Average T_ao_ (time for 80% decay from [DA]_o_ peak) in the NAc. Adults-NR (black, n = 8), Adults-PR (blue, n = 7), Adolescents-NR (grey, n = 9) and Adolescents-PR (orange, n = 5). Bars show means ± SEM and circles show individual (*e*.*g*. recording site) data points. * *p* < 0.05 Diet effect (Student’s unpaired t-test), # *p* < 0.05 Age effect (two-way ANOVA).

#### Frequency-dependent NAc dopamine release

Dopamine neurons *in vivo* show a range of responses from low-frequency firing (< 10 Hz, *tonic* mode) to brief bursts of action potentials at high frequency (15-25 Hz, *phasic* mode) [59–62]. We therefore investigated the effect of protein restriction on dopamine release at different stimulation frequencies ranging from 1 to 20 Hz (1 p = 1 Hz, or 5 p at 5, 10 or 20 Hz). Evoked dopamine release increased with the stimulation frequency (**Figure 3A-B**; three-way repeated measures ANOVA: Frequency, F(3,72) = 39.2, *p* < 0.001) similarly in adolescent and adult groups (Age, F(1,24) = 3.1, *p* = 0.09). Protein restriction did not affect the frequency-dependent effect on dopamine release (Frequency x Diet, F(3,72) = 1.6, *p* = 0.2). However, protein restriction did differentially affect NAc dopamine release depending on age (Diet, 228 F(1,24) = 1.8, *p* = 0.2; Age x Diet, F(1,24) = 13.4, *p* < 0.01; Frequency x Diet x Age, F(3,72) = 229 3.2, *p* < 0.05).

**Figure 3.**
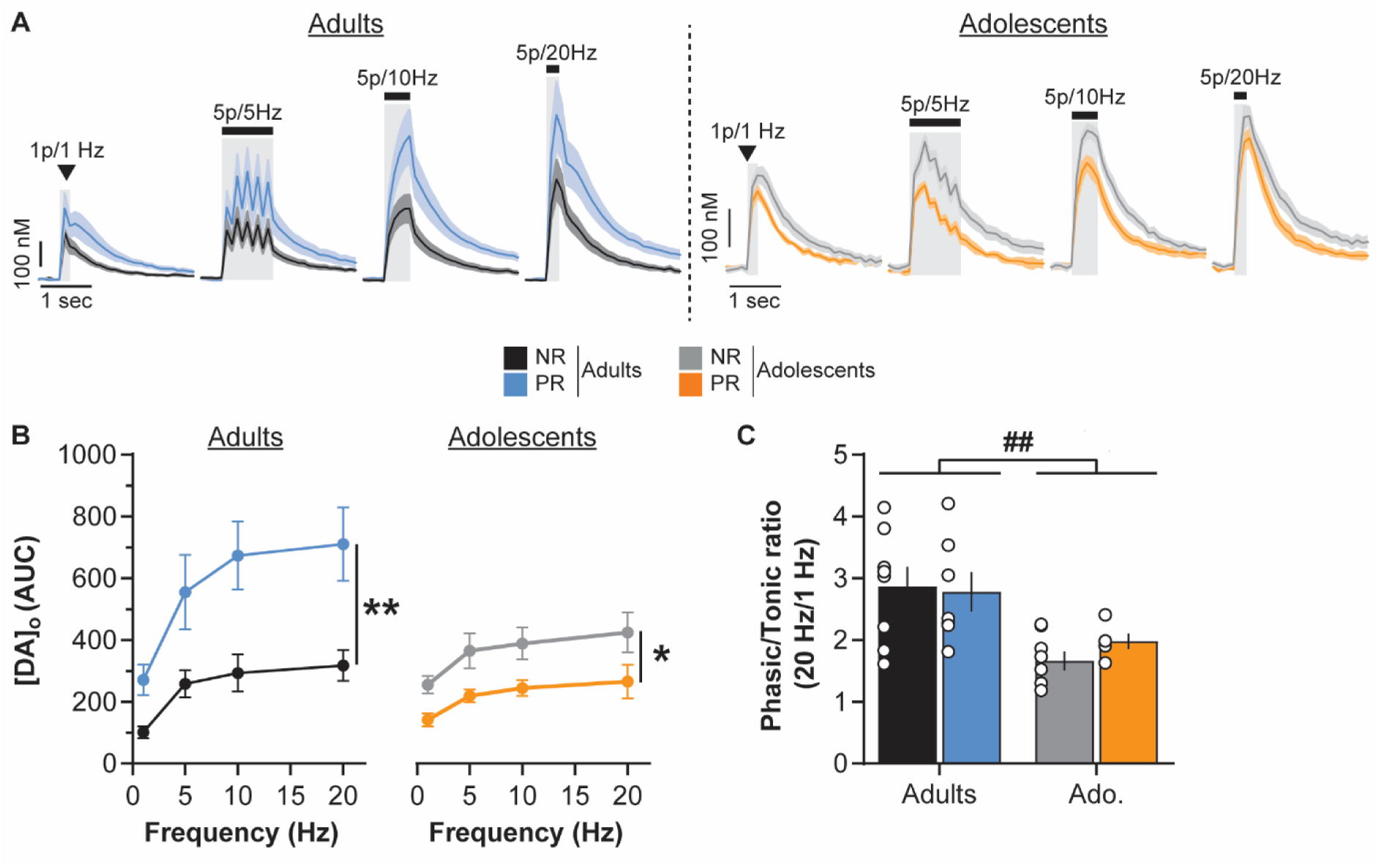
Age-dependent impact of protein restriction on frequency-dependent NAc dopamine release. (A) NAc [DA]_o_ versus time (in nM, mean ± SEM) for each diet group (non-restricted, NR; protein-restricted, PR) aligned to the electrical stimulation (black symbol) at 1 Hz (single pulse), 5, 10 or 20 Hz (5 pulses; 0,7 mA, 0.2 ms). (B) Protein restriction increased frequency dependent NAc dopamine release (AUC) in adult rats (*left*) but decreased it in adolescent rats (*right*). (C) Protein restriction has no impact on NAc [DA]_o_ phasic/tonic ratios. Adults-NR (black, n = 8), Adults-PR (blue, n = 7), Adolescents-NR (grey, n = 9) and Adolescents-PR (orange, n = 5). Bars show means ± SEM and circles show individual (e.g. recording site) data points. *\* p <* 0.05 Diet effect (two-way ANOVA followed by Sidak’s *post hoc* tests), ## *p* < 0.01 Age effect (two-way ANOVA).

In adult rats, protein restriction increased dopamine release in response to the range of stimulation frequencies (two-way repeated measures ANOVA: Diet, F(1,13) = 9.3, *p* < 0.01; 233 Frequency, F(3,39) = 43.6, *p* < 0.001; Diet x Frequency, F(3,39) = 5.3, *p* < 0.01; Sidak’s *post hoc* tests: all *p* < 0.05). Conversely, protein restriction during adolescence significantly decreased evoked NAc dopamine release (two-way repeated measures ANOVA: Diet, F(1,11)= 6.1, *p* < 0.05; Frequency, F(3,33) = 6.3, *p* < 0.01; Diet x Frequency, F(3, 33) = 0.1, *p* = 0.9).

The relationship between dopamine release during tonic and phasic activity is a central process in the signaling of significant environmental events and learning [62,65,66]. We examined whether protein restriction during either adolescence or adulthood affected the ‘phasic/tonic ratio’ of NAc dopamine release (5 p at 20 Hz / 1 p, **Figure 3C**). Adolescent rats exhibited a lower ratio than adult rats (Two-way ANOVA: Age, F(1,24) = 13.9, *p* < 0.01). However, protein restriction did not alter this ratio at either age (Diet, F(1,24) = 0.2, *p* = 0.7; 243 Age x Diet, F(1,24) = 0.6, *p* = 0.5).

### Age dependent impact of protein restriction on dorsal striatum dopamine release

#### Single pulse evoked striatal dopamine release

In the dorsal striatum, protein restriction had no effect on dopamine release evoked by single pulse stimulation whether rats were exposed to the diet during adulthood or adolescence 250 (**Figure 4A-B**; two-way ANOVA: Age, F(1,28) = 0.7, *p* = 0.4; Diet, F(1,28) = 0.01, p = 0.9; Diet x Age, F(1,28) = 1.3, p = 0.3), which is also confirmed by the distribution analysis (**Figure 4C**; Kolmogorov-Smirnov tests: Adult groups, D(14) = 0.3, p = 0.8; Adolescent groups, D(14) = 0.6, p = 0.1). Moreover, protein restriction also did not significantly affect [DA]_o_ peak amplitude 254 (**Figure 4D**; Two-way ANOVA: Diet, F(1,28) = 0.04, p = 0.9; Age, F(1,28) = 0.08, p = 0.8, Diet 255 x Age, F(1,28) = 1.4, p = 0.2). or dopamine clearance (**Figure 4E**; Two-way ANOVA: Diet, 256 F(1,28) = 0.03, p = 0.9; Age, F(1,28) = 0.9, p = 0.4, Diet x Age, F(1,28) = 2.1, p = 0.2; see also **Supplementary Figure 1**).

**Figure 4.**
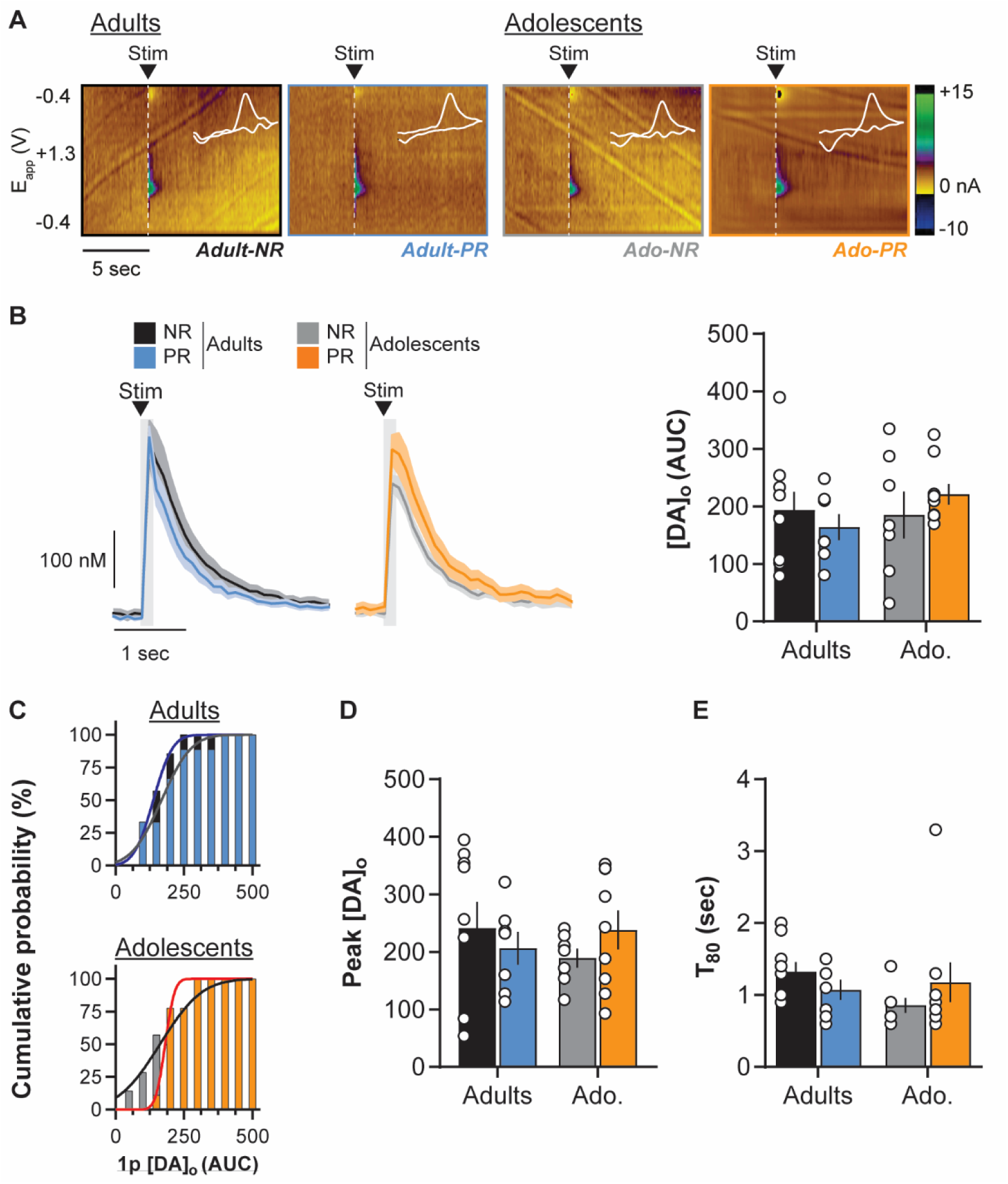
Age-dependent impact of protein restriction on dorsal striatum dopamine release evoked by single pulse. (A) Representative FSCV color plots for each diet group (non-restricted, NR; protein-restricted, PR) depicting current changes (color) over time (x-axis; in sec) as a function of the recording electrode holding potential (y-axis; −0.4 to +1.3 V and back) in response to single pulse electrical stimulation (0.7 mA, 0.2 ms; vertical white dashed lines). White line insets represent voltammograms for each color plot. (B) **Left**: Dorsal striatum [DA]_o_ (in nM; mean ± SEM) in slices from adult and adolescent NR and PR rats, aligned to the single pulse electrical stimulation (black arrow); **Right** [DA]_o_ release (AUC) evoked by single pulse stimulation in the dorsal striatum. (C) Cumulative distribution of single pulse evoked dorsal striatum [DA]_o_AUC in adult **(top)** and adolescent **(bottom)** groups. (D) Mean [DA]_o_ peak evoked by single pulse stimulation in the dorsal striatum. (E) Average T_80_ (time for 80% decay from [DA]_o_ peak) in the dorsal striatum. Adults-NR (black, n = 9), Adults-PR (blue, n = 7), Adolescents-NR (grey, n = 7) and Adolescents-PR (orange, n = 9). Bars show means ± SEM and circles show individual (*e*.*g*. recording site) data points.

#### Frequency-dependent striatal dopamine release

As previously observed in the NAc, striatal dopamine release increased as a function of the stimulation frequency (**Figure 5A-B**; three-way repeated measures ANOVA: Frequency, F(3,84) = 13.06, p < 0.001) similarly in both age groups (Age, F(1,28) = 1.5, *p* = 0.2; Frequency x Age, F(3, 84) = 2.1, *p* = 0.1). However, first analyses suggested that protein restriction did not significantly change dopamine release evoked by all frequencies tested (Diet, F(1,28) = 265 3.5, *p* =0.07; Diet x Frequency, F(3,84) = 1.8, *p* = 0.1; Diet x Age, F(1,28) = 2.6, *p* = 0.1; Diet 266 x Frequency x Age, F(3,84) = 1.9, *p* = 0.1).

**Figure 5.**
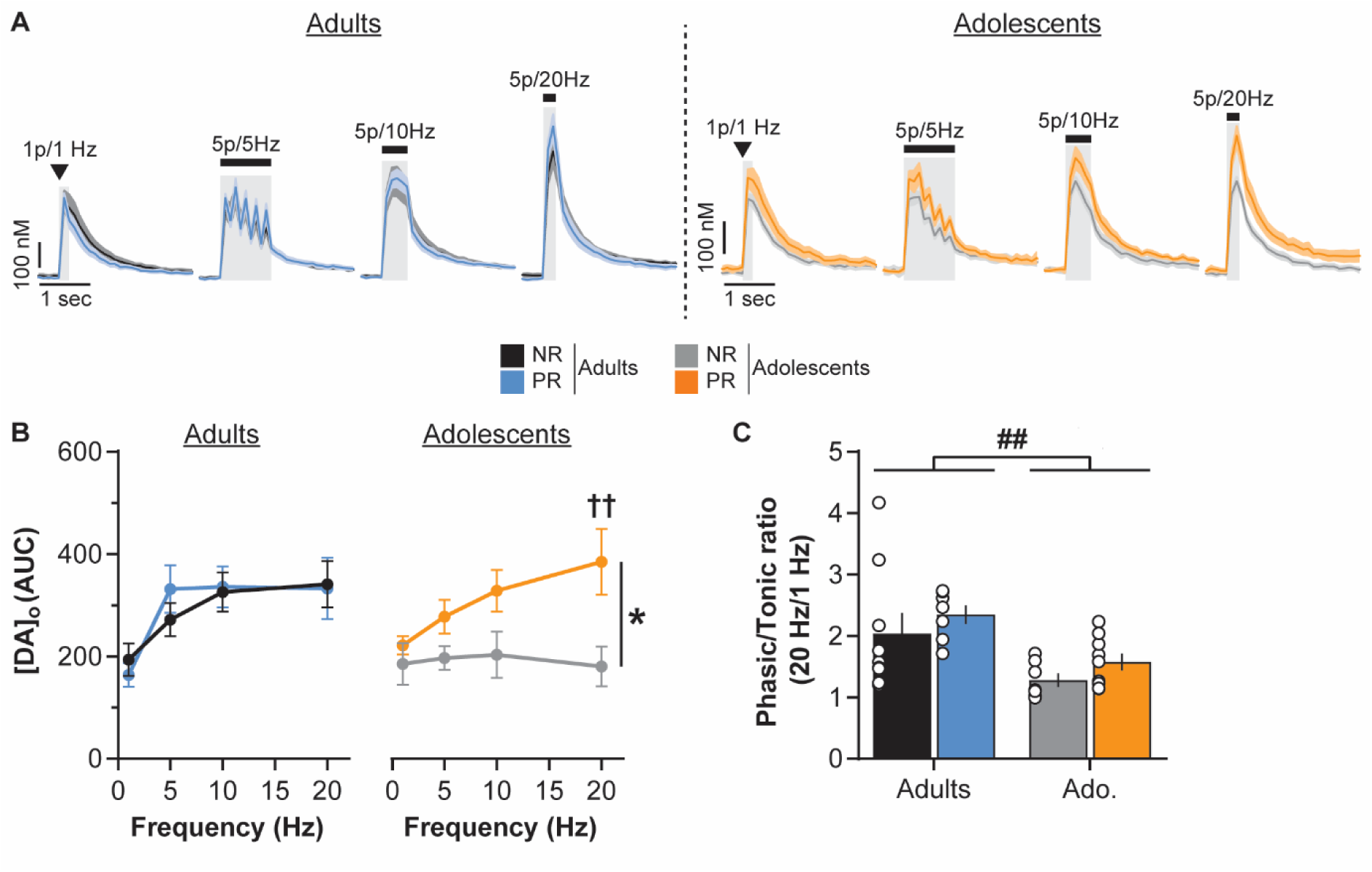
Age-dependent impact of protein restriction on frequency-dependent dorsal striatum dopamine release. (A) Dorsal striatum [DA]_o_ (in nM, mean ± SEM) for each diet group (non-restricted, NR; protein-restricted, PR) aligned to the electrical stimulation (black symbol) at 1 Hz (single pulse), 5, 10 or 20 Hz (5 pulses; 0.7 mA, 0.2 ms). (B) Protein restriction at adulthood did not affect dorsal striatum dopamine release (AUC) in adults (*left*) but increased it in adolescent rats (*right*). (C) Protein restriction has no impact on [DA]_o_ phasic/tonic ratios. Adults-NR (black, n = 9), Adults-PR (blue, n = 7), Adolescents-NR (grey, n = 7) and Adolescents-PR (orange, n = 9). Bars show means ± SEM and circles show individual (*e*.*g*. recording site) data points. * *p* < 0.05 Diet effect (two-way ANOVA followed by Sidak’s *post hoc* tests), †† *P* < 0 01 Frequency effect (two-way ANOVA followed by Dunnett’s *post hoc* tests versus 1 Hz), ## *p* < 0.01 Age effect (two-way ANOVA).

Separate analyses for each age group confirmed that protein restriction has no significant effect on frequency-dependent striatal dopamine release in adults (two-way repeated 270 measures ANOVA: Diet, F(1,14) = 0.02, *p* = 0.9; Frequency, F(3,42) = 26.6, *p* < 0.001; Diet x Frequency, F(3, 42) = 1.8, *p* = 0.2). In contrast, protein restriction in adolescent rats significantly increased stimulation-evoked striatal dopamine release (two-way repeated measures ANOVA: Diet, F(1,14) = 8.7, *p* < 0.05; Frequency, F(3,42) = 1.8, *p* = 0.2; Diet x Frequency, F(3, 42) = 1.9, *p* = 0.1), especially in response to phasic-like stimulations (Sidak’s post hoc tests: 1-10 Hz all *p* > 0.1; 20 Hz *p* < 0.01). Moreover, the frequency-dependent increase in dorsal striatum dopamine release is significantly observed in the adolescent PR group (Dunnett’s *post hoc* tests versus 1 Hz stimulation: 5 Hz, *p* = 0.5; 10 Hz, *p* = 0.08; 20 Hz, *p* < 0.01) but not in the NR control group (5-20 Hz versus 1 Hz stimulation, all *p* > 0.9). This last result suggests that the nigrostriatal dopamine system may be sensitized by protein restriction during adolescence, despite an overall decrease in evoked release of dopamine.

Similar to what we observed in the NAc, the ‘phasic/tonic’ ratio of striatal dopamine release was lower in adolescent slices (**Figure 5C**; Two-way ANOVA: Age, F(1,28) = 11.7, p < 0.01) but was not altered by protein restriction (Diet, F(1,28) = 1.8, p = 0.2; Age x Diet, F(1,28) = 284 0.001, p = 1.0).

## DISCUSSION

Protein homeostasis is a crucial physiological function for almost all species throughout the lifespan. Despite the deleterious consequences of protein restriction on a multitude of physiological functions, the neurobiological impact of such a diet at different ages remains largely unexplored. The present study reveals that protein restriction affects the function of the mesolimbic and nigrostriatal dopamine pathways. More importantly, our results demonstrate that these effects are dependent on the age at which protein restriction is experienced, highlighting adolescence as a vulnerability window for the deleterious effects of an unbalanced diet.

The impact of protein restriction on weight is highly dependent on the degree of restriction and the physiological state of the animal [9,10,55]. When performed at adulthood, protein restriction did not affect rats’ weight, consistent with our previous results [11]. Moreover, adult rats slightly increased their daily energy intake relative to their body weight. In adults, this increase may explain the absence of effect on weight as rats attempt to compensate protein deficiency with a general hyperphagia [11]. An alternative explanation is that low protein diet may change energy expenditure, as previously observed [67]. In contrast, protein restriction during adolescence significantly limits animals’ normal trajectory of weight gain. As for adults, adolescent PR rats increased their daily energy intake compared to the control NR group. Adolescent animals are rapidly growing and have higher protein requirements than adults [55]. Surprisingly, this change in food intake behavior did not seem to be sufficient to support normal growth. In the present study, the low protein diet (5% protein from casein) was the only source of nutrients. Breakdown analysis of macronutrient intake revealed that the important protein deficiency observed in PR groups is associated with an indirect increase in carbohydrate and fat intake contained in animal food. The regulation of protein appetite and the balance between protein intake and other macronutrients is still poorly understood but several studies suggest that numerous species regulate their food-related behaviors to avoid protein deficiency [8–10], which may lead to the overconsumption of other nutrients. It remains intriguing, however, that in this case adolescent PR rats did not exhibit a larger increase of their food intake. As both the overconsumption of sweet or fat diets may impact the functioning of the dopamine system especially during development [43–47], we cannot exclude that the diet impact reported here may be the result of the combination of protein deficiency and concurrent changes in carbohydrate and fat intake.

The two main dopamine projections to the NAc and the dorsal striatum are involved in various food-related processes including incentive salience [16] and prediction error [66], using taste and nutritional (post-ingestive) values of food [23–28]. Here, we observed that protein restriction differentially affected projection-specific dopamine release depending on age of diet exposure. At adulthood, protein restriction increased NAc dopamine release but had no effect on dorsal striatum dopamine release. Tonic and phasic dopamine firing and release convey different information about motivational and learning processes [16,19,23,66,68]. In the mesolimbic pathway, PR diet at adulthood increased both responses to low ‘tonic’ and high ‘phasic’ stimulations but did not alter the phasic/tonic ratio, suggesting a more general increase in the capacity of terminals for dopamine release rather than an change in the contrast between different dopamine signaling modes [59,69]. Such global sensitization of the mesolimbic pathway may profoundly alter motivated behaviors like food preferences [11,13], and increase the rewarding properties of protein-enriched food in restricted/deprived animals 331 [12].

Protein restriction during adolescence had a broader impact on the function of dopamine terminals, relative to the same diet during adulthood. In contrast to what we observed at adulthood, protein restriction in adolescents decreased NAc dopamine release both in response to single pulse stimulation, low frequency pulse trains (5-10 Hz) and high frequency burst-like stimulation (5p at 20 Hz). Dopamine neurons exhibit an elevated firing rate during adolescence [50,53,54] associated with changes in dopamine availability in dopamine projection targets [48,49,51]. Based on this and our first results showing an effect of protein restriction at adulthood on NAc dopamine release, we might have expected an enhancement of the diet effect during adolescence. One way to reconcile these opposite findings is to consider that the degree of protein restriction in adolescent rats may be more profound than in adults. As discussed earlier, we observed a substantial impact of protein restriction on weight gain in protein-restricted adolescents (and not in adults) suggesting a more severe level of restriction. As dietary protein is a major source of amino acids (e.g. tyrosine) required for catecholaminergic metabolism (synthesis, release, enzymatic activity), one hypothesis is that a greater protein deficiency in adolescent rats than adults will affect average dopamine levels and the ability to synthesize and release dopamine. Accordingly, previous studies have reported a decrease in dopamine in several brain regions in response to pre- or perinatal protein malnutrition as well as an hypo-responsivity to psychostimulants (see [5] for review). In the dorsal striatum in adolescents, we observed an opposite pattern compared to the NAc. As such, evoked dopamine release was increased after adolescent protein restriction, especially at high stimulation frequencies. Such an effect partially rules out the hypothesis of a global amino acid deficiency. However, the nigrostriatal dopamine pathway matures earlier than other dopamine pathways [48] and may then be less sensitive to protein restriction. Striatal and NAc dopamine pathways are involved in different aspects of food-related processes and recent advances demonstrated that striatal, but not NAc, dopamine signaling is involved in encoding the nutritional value of food [70]. The increase in evoked dopamine release in striatal areas only seen in adolescent-exposed rats reported in the present study may support a nutrition-seeking response to the elevated protein requirement at this age.

The effect of protein restriction at adulthood or during adolescence on dopamine pathways may also involve regulation of dopamine terminal activity by reuptake processes or local striatal microcircuits [65,69]. Dopamine reuptake activity may be changed by specific diets [39,40]. Here, we did not observe any significant change induced by protein restriction on dopamine clearance in response to single pulse stimulations. Combined with the absence of significant diet effects on the [DA]_0_ peak amplitudes, this suggests that neither protein restriction during adolescence nor adulthood impacts dopamine transporter functioning. However, we cannot totally exclude reuptake changes as we observed diet-dependent changes in evoked dopamine release quantified by AUC. The AUC could vary because of changes in either dopamine release or reuptake. On the other hand, striatal microcircuits also mature during adolescence [75,76] and may be sensitive to different diet effects. These issues and the behavioral consequences of dietary protein alterations on the dopamine system remain to be investigated.

The direct influence of protein or amino acids levels on dopamine neurons is still unexplored, however, these neurons receive input from hypothalamic regions which are able to detect amino acids [10,14]. Protein restriction also induces a broad metabolic response involving peripheral food-related signals to which dopamine neurons are directly sensitive [31–35]. Dopamine release is especially sensitive to insulin through its actions at specific receptors located both directly on dopamine neurons [77] and on striatal cholinergic neurons [37]. The effects of insulin on the dopamine system and dopamine-related behaviors are complex and depend on insulin concentration, brain region, cell type and the current physiological state [40,78]. Protein restriction is known to increase insulin sensitivity and glucose metabolism [13,79], which may then modulate dopamine’s neurobiological and behavioral functions. The interaction of the dopamine and insulin systems in response to different diets differing in protein content warrants further *ex vivo* and *in vivo* investigation.

In conclusion, our study provides evidence that prolonged protein restriction has an important impact on function of dopamine terminals in the NAc and dorsal striatum. More importantly we highlight the increased sensitivity of the dopamine system during adolescence to the deleterious effects of a diet that is inadequate in protein. Adolescence is characterized by important maturation events within dopamine circuitry and dopamine-related processes [48– 52,54] and numerous studies have now demonstrated that adolescence is an important vulnerability window for diet-related alterations of cognitive and neurobiological functions [43– 47]. How protein restriction during adolescence may have different, and potentially long-term, impacts on dopamine-related behaviors considering its opposite effects on the mesolimbic and nigrostriatal pathways, remains to be investigated. Given the role of malnutrition and inadequate protein intake on neurodevelopmental psychiatric disorders [5,6] involving alterations of the dopamine system [17,80,81] and having their onset during adolescence [36,82], our current findings also represent a step towards a better understanding of the mechanisms regulating protein appetite, protein malnutrition, and the emergence of dopamine-related disorders.

## ACKNOWLEDGEMENTS

The authors would like to acknowledge the help and support from the staff of the Division of Biomedical Services, Preclinical Research Facility, University of Leicester, for technical support and the care of experimental animals.

## AUTHOR CONTRIBUTIONS

FN, KZP and JEM designed research; FN performed research, FN, KZP, AMJY and JEM analyzed data; FN, KZP, AMJY and JEM wrote the manuscript.

## SUPPLEMENTARY INFORMATION

**Supplementary Table 1.**
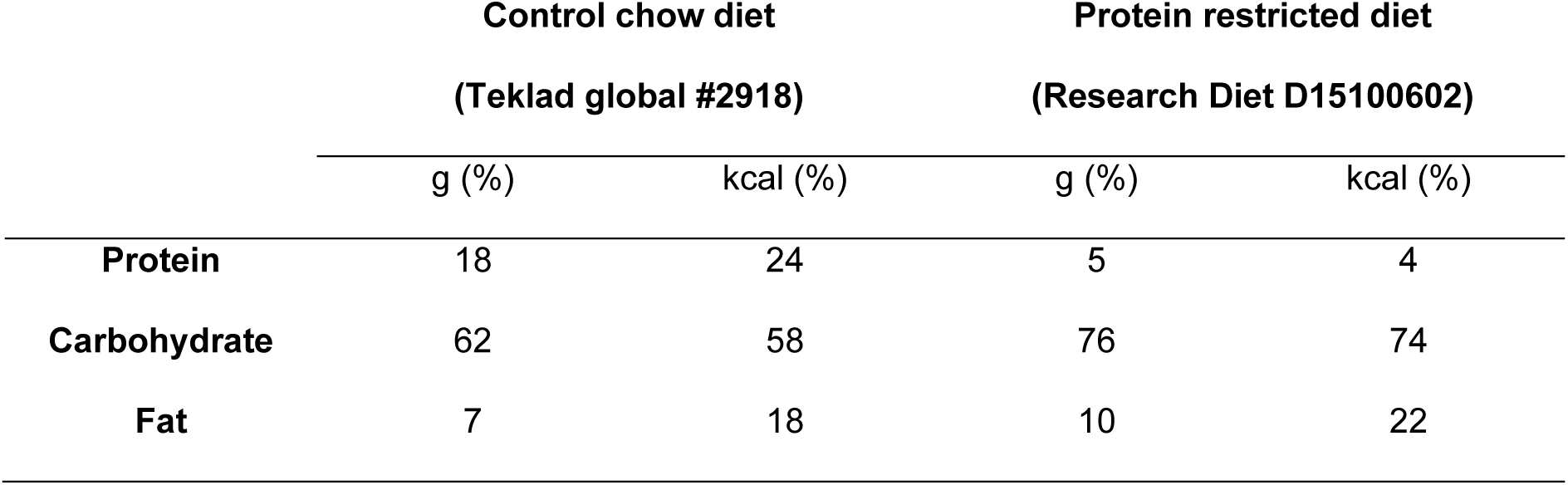
Macronutrient composition of the control chow and protein restricted diet.

**Supplementary Table 2.**
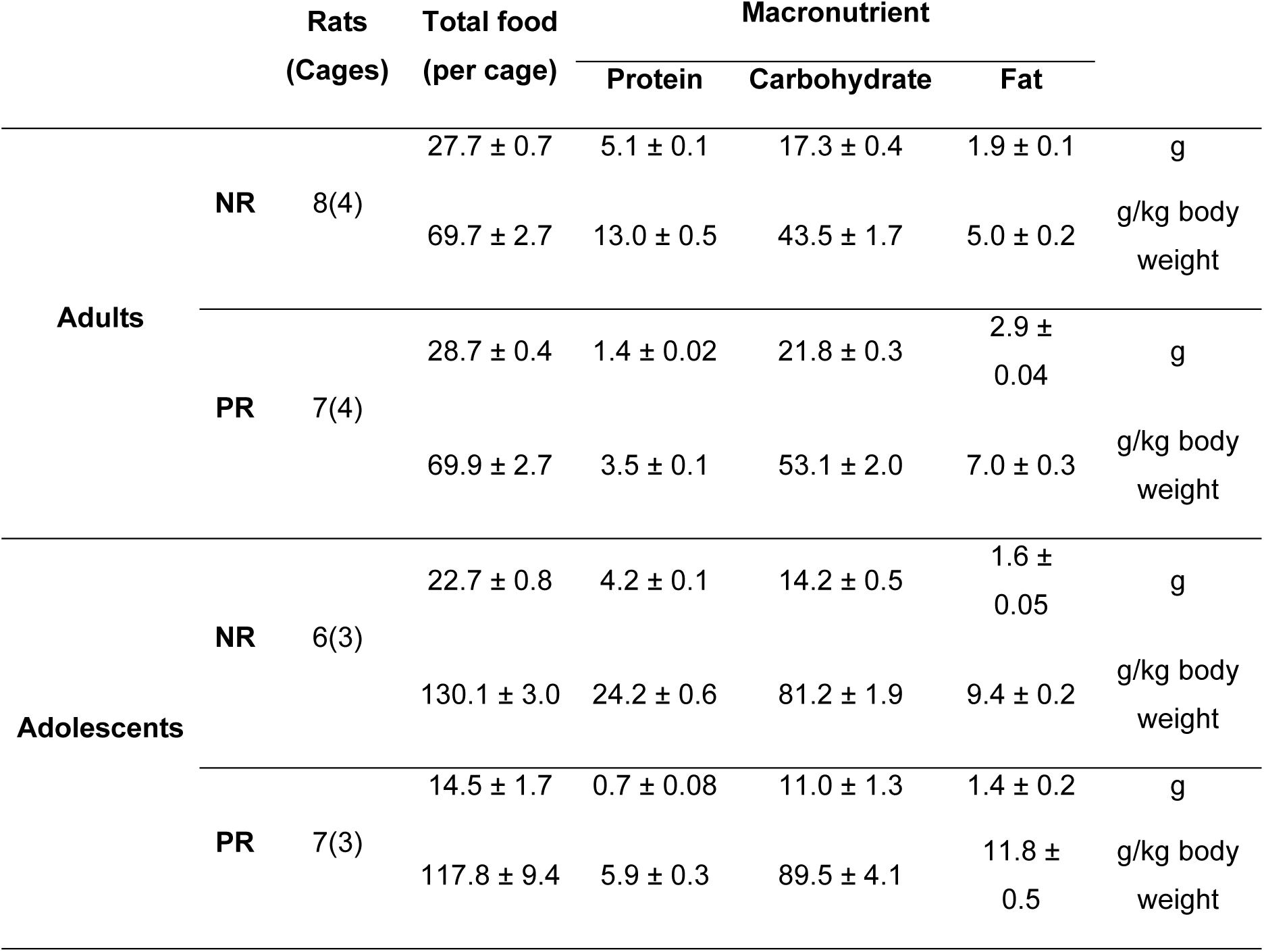
Macronutrient breakdown of average daily food intake per cage (in g) or normalized to rats’ body weight (g/kg body weight) for each diet condition.

**Supplementary Figure 1.**
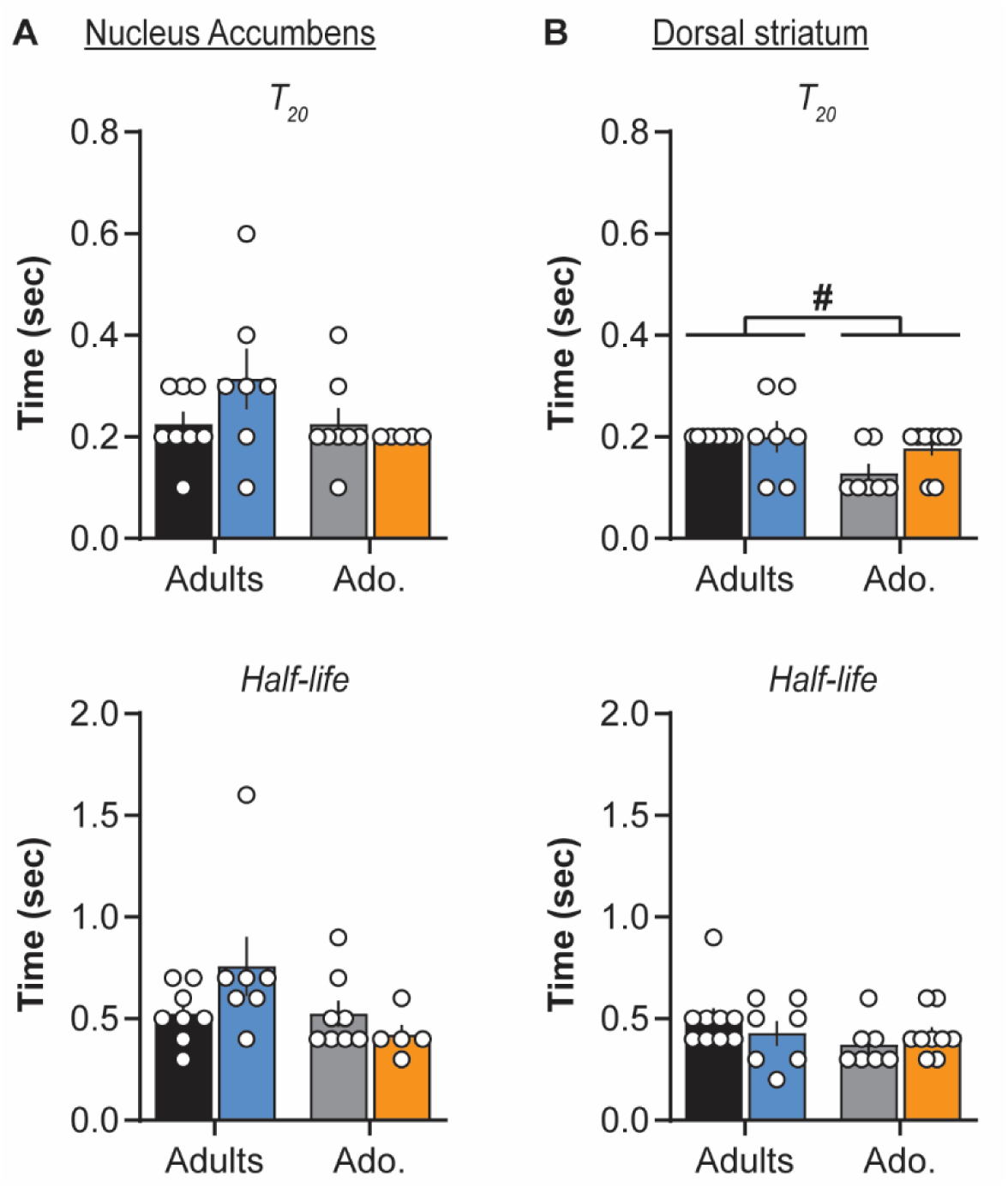
Protein restriction at adulthood or during adolescence did not change dopamine clearance parameters (T_20_: time for 20% decay from [DA]_o_ peak; Half-life: time for 50% decay from [DA]_o_ peak) after single pulse stimulation in the NAc (A; T_20_ Age F(1,24) = 2.2, *p* = 0.2; Diet F(1,24) = 0.7, *p* = 0.4; Age **x** Diet F(1,24) = 2.2, *p* = 0.2 / Half-life Age F(1,24) = 3.5, *p* = 0.07; Diet F(1,24) = 0.5, *p* **-** 0.5; Age **x** Diet F(1,24) = 3.5, *p* **-** 0.07) or in the dorsal striatum (B; T_20_ Age F(1,28) = 6.9, *p* < 0.05; Diet F(1,28) = 1.9, *p* = 0.2; Age **x** Diet F(1,28) = 1.9, *p* = 0.2 / Half-life Age F(1,**28**) = 1.9, *p* = 0.2; Diet F(1,28) = 0.04, *p* = 0.8; Age **x** Diet F(1,28) = 1.6, *p* = 0.2). Adults-NR (black), Adults-PR (blue), Adolescents-NR (grey) and Adolescents-PR (orange). Bars show mean ± SEM and circles show individual (*e*.*g*. recording site) data points. # *p* < 0.05 Age effect (two-way ANOVA)

